# Metabolic modeling of *Streptococcus mutans* reveals complex nutrient requirements of an oral pathogen

**DOI:** 10.1101/419507

**Authors:** Kenan Jijakli, Paul A. Jensen

## Abstract

*Streptococcus mutans* is a Gram positive bacterium that thrives under acidic conditions and is a primary cause of tooth decay (dental caries). To better understand the metabolism of *S. mutans* on a systematic level, we manually constructed a genome-scale metabolic model of the *S. mutans* type strain UA159. The model, called iSMU, contains 656 reactions involving 514 metabolites and the products of 488 genes.

We interrogated *S. mutans*’ nutrient requirements using model simulations and nutrient removal experiments in defined media. The iSMU model matched experimental results in greater than 90% of the conditions tested. We also simulated effects of single gene deletions. The model’s predictions agreed with 78.1% and 84.4% of the gene essentiality predictions from two experimental datasets. Our manually curated model is more accurate than *S. mutans* models generated from automated reconstruction pipelines. We believe the iSMU model is an important resource for understanding how metabolism enables the cariogenicity of *S. mutans*.

## Introduction

*Streptococcus mutans* is one of over 600 species of bacteria in the oral microbiome. This Gram-positive, lactic acid bacterium thrives in the oral environment in part due to its metabolic flexibility. *S. mutans* can feed on several carbohydrates (1) and has complex, interdependent amino acid auxotrophies (2). *S. mutans* is the primary cause of tooth decay (dental caries). By fermenting a wide array of dietary sugars into lactic acid, *S. mutans* creates a highly acidic microenvironment near the tooth surface (as low as pH 3.0). The lactic acid demineralizes the tooth structure, resulting in decay.

Understanding the acidogenic capabilities of the *S. mutans* requires an unbiased, systems-level approach. Previous studies have shown that acid production and tolerance in *S. mutans* requires large changes in gene expression and metabolic pathway utilization (3). For example, decreasing pH increases glycolytic activity and branched-chain amino acid synthesis without increasing cell growth (4). A drop in pH is also accompanied by an increased expression of F-ATPases to maintain a higher intracellular pH (5).

Mathematical models aid in our understanding of how an organism’s genes collectively give rise to a phenotype. Models translate bioinformatic features (differential expression, presence/absence of genes) into biological function (flux distributions, uptake and secretion rates, and fitness). Constraint-based reconstruction and analysis (COBRA) of genome-scale models is widely used to integrate genetic and metabolic data to produce phenotypic predictions (6, 7). Models of microbial metabolism and transcriptional regulation predict responses to gene deletions (8, 9), mutation (10, 11), metabolic shifts (12, 13), and long-term evolution (14, 15). The models identify emergent properties of a metabolic network, including links between pathways and inter-dependencies among genes (16, 17).

We present iSMU v1.0, a genome scale metabolic model for the *S. mutans* type strain UA159. Our model is manually curated using multiple databases, literature evidence, and phenotyping experiments. Our investigation of *S. mutans* focused our metabolism for two reasons: 1.) the primary metabolic products of *S. mutans*, lactic acid and biofilm matrix, are responsible for the pathogen’s cariogenicity; and 2.) metabolic networks are among the best characterized intracellular networks with established computational techniques. Given metabolism’s central role in cariogenesis, we believe the iSMU model will improve our understanding *S. mutans*’ role in oral health.

## Materials and Methods

### Model Construction

The metabolic network of *S. mutans* UA159 was reconstructed following best practices in the COBRA modeling community (18). As summarized in Figure 1A, reconstruction began with the annotated UA159 genome (RefSeq GCA_000007465.2). Metabolic enzymes and the associated reactions were initially collected from KEGG (19) and Uniprot (20). The Metacyc (21), RHEA (22), ModelSEED (23), BiGG (24), and ChEBI (25) databases were used as secondary sources for metabolic reactions. Transport reactions were verified with TransportDB (26). When possible, KEGG identifiers were used for metabolites and reactions for consistency with other databases. Custom identifiers (e.g. “add00001”) were added for reactions or metabolites without KEGG identifiers. Reactions without gene associations were only added when supported by experimental or literature evidence. These 11 reactions are explained in Table S1. All chemical species and formulas were converted to their protonation state at pH 7.0 using the ModelSEED database. A custom map of the iSMU model was constructed using Escher version 1.6.0 (Figure 2) (27).

**Figure 1:**
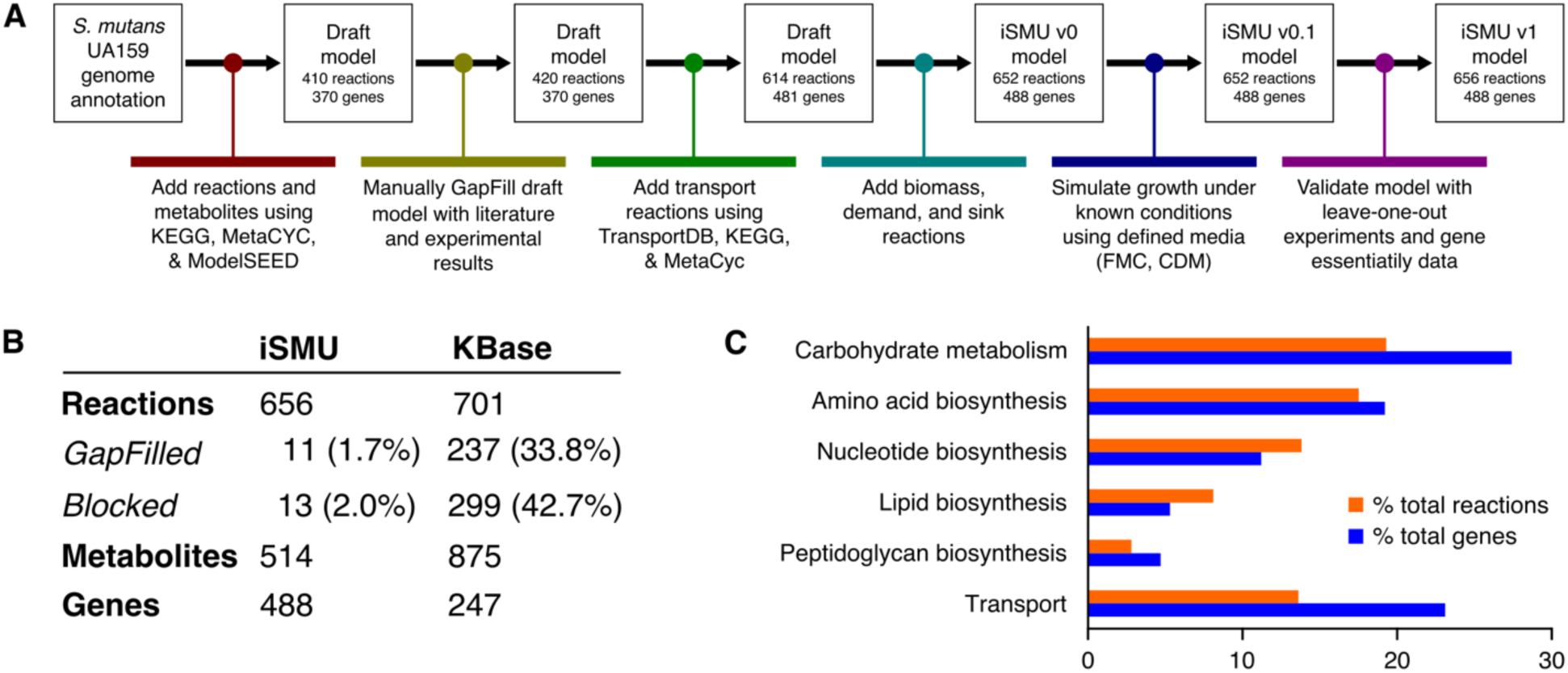
**A.** Reconstruction of the *S. mutans* metabolic network began with an annotated UA159 genome. The draft model was refined with bioinformatics databases and experimental results. **B.** The manually reconstructed model (iSMU) has fewer GapFilled (non-gene associated) and blocked reactions than a model generated automatically by the KBase systems. **C.** The iSMU model contains reactions across multiple KEGG pathways, especially carbohydrate catabolism and transport.

**Figure 2:**
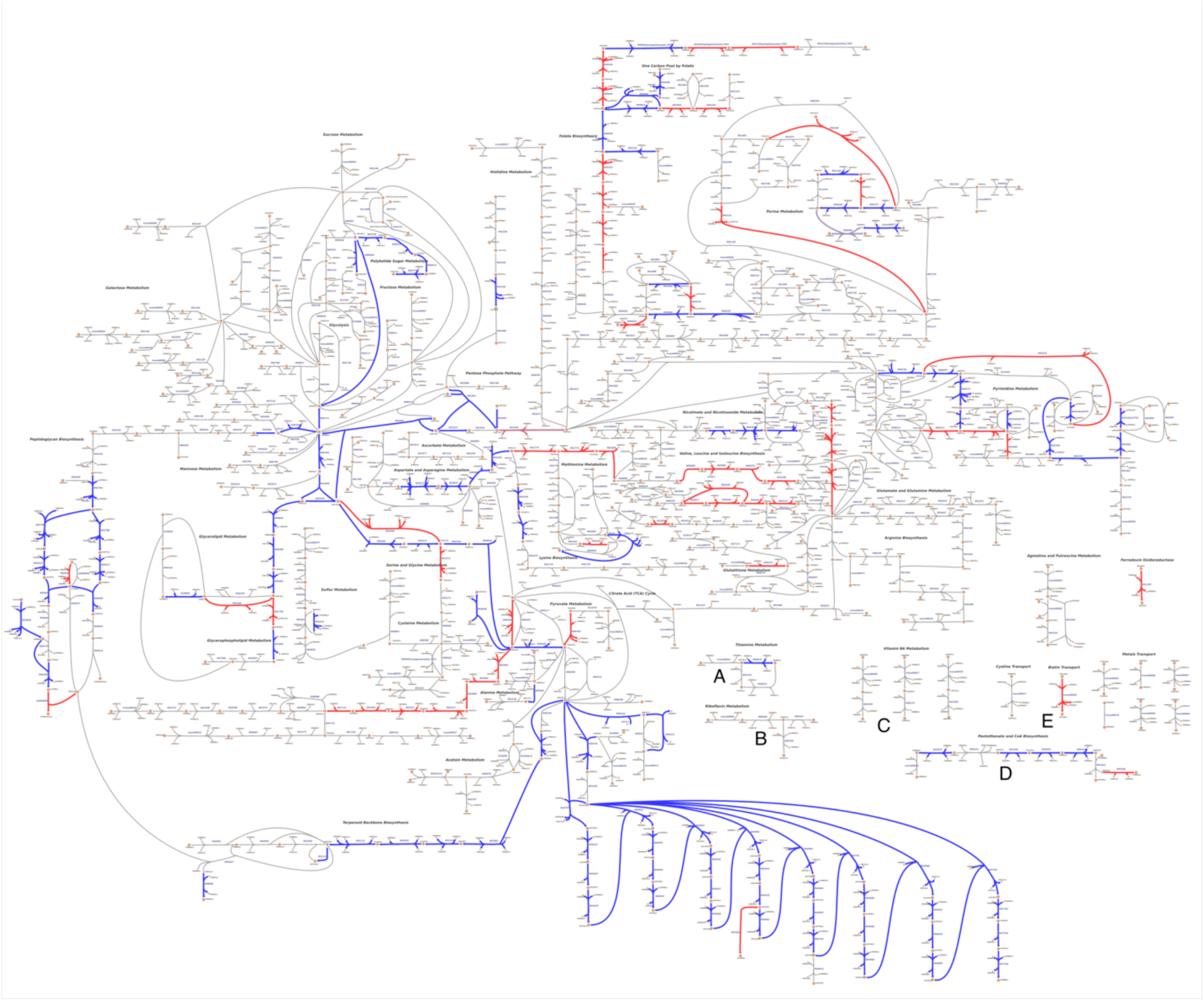
A custom pathway map shows all reactions in the iSMU model. (A high-resolution image is available as File S4.) *S. mutans* UA159 appears to lack complete pathways for synthesizing thiamine (**A**), riboflavin (**B**), pyridoxine (**C**), pantothenate (**D**), and biotin (**E**). Reactions are colored by agreement between the essentiality predictions of the associated genes and Tn-seq data from Shields et al. [ref]. Blue reactions agree with the Tn-seq essentiality results; red reactions disagree. Overall agreement between the datasets is 84.8% (see Figure 4).

### Model Simulations using Flux Balance Analysis

To simulate growth using iSMU, reactions were collected into a stoichiometric matrix *s* where element *s*(*i*, *j*) corresponds to the stoichiometric coefficient of the model’s^th^ metabolite in the *j*^th^ reaction. *s*(*i*, *j*) is negative if the metabolite is consumed and positive if the metabolite is produced. Two vectors of lower (*l*) and upper (*u*) bounds determine the reversibility of reactions. A vector of reaction fluxes ν was calculated by maximizing the flux through the biomass reactions subject to mass balance constraints (*sν* = 0) and reversibility constraints (*l* ≤ ν ≤*u*). To simulate gene deletions, the gene/protein/reaction rules for each reaction were evaluated to identify reactions that cannot carry flux in the deletion strain. The upper and lower bounds of these reaction were set to zero before maximizing flux through the biomass reaction. Genes were considered essential if their deletion allowed no biomass flux.

All simulations were performed with Matlab (version R2016b; MathWorks [https://www.mathworks.com]) using the COBRA toolbox (28). Mathematical programs were solved with Gurobi Optimizer (version 7.5; Gurobi Optimization [http://www.gurobi.com]). Gene set enrichment for KEGG pathways was performed using DAVID (29, 30).

### Model Availability

The final model is available as an SBML file and a spreadsheet compatible with the COBRA toolbox (Files S1 & S2). The model map is available as a JSON file (File S3) and SVG image (File S4). Future versions of the model and map will be available on the authors’ website [http://jensenlab.net].

### Strains and Media

*S. mutans* UA159 (ATCC 700610) was cultured on Brain-Heart Infusion (Sigma) agar plates or in Todd Hewitt broth with 0.3% yeast extract (Sigma). Strains were grown overnight in 5% CO_2_ at 37°C unless specified.

### Growth Assays

Growth experiments were performed in a Chemically Defined Medium (CDM) (31) with 22 amino acids, 11 vitamins, 3 nucleobases, 8 inorganic salts, and glucose (Table 1). Complete CDM or leave-one-out variants were prepared fresh weekly from concentrated stocks (31). All components were purchased from Sigma-Aldrich USA and were sterilized by autoclaving or filtration.

**Table 1:**
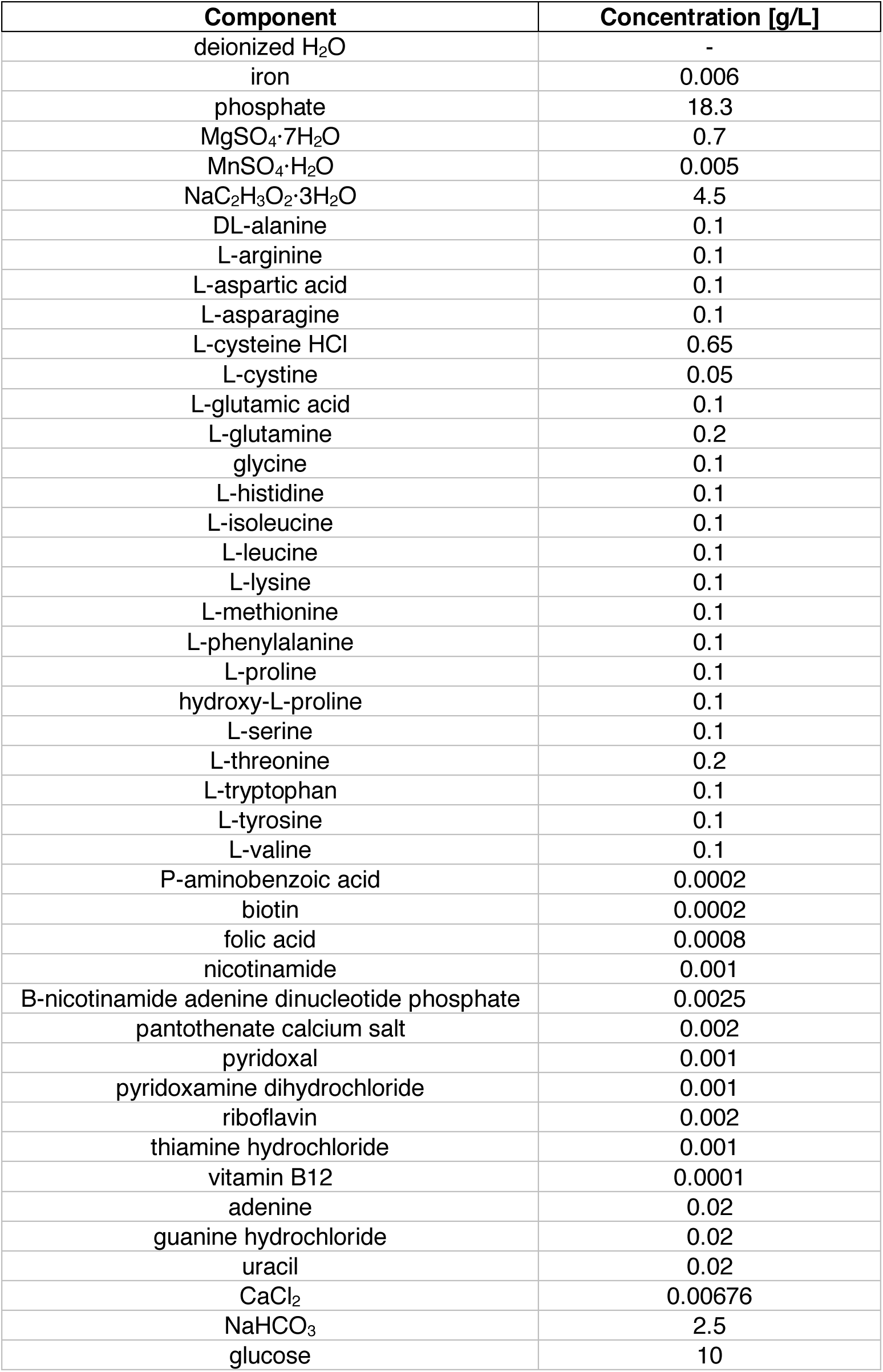
CDM is composed of 22 amino acids, 11 vitamins, 3 nucleobases, 8 inorganic salts, and glucose.

Overnight cultures of *S. mutans* were washed three times in sterile water. The overnight culture was concentrated 5 × (from 5 ml to 1 ml), and 1 μl of the concentrate was used to inoculate wells of a 96 well plate containing 200 μl of defined media. Plates were incubated without agitation in 5% CO_2_ at 37°C. Optical density was measured by absorbance at 600 nm every hour for 16 h using a Tecan Infinite 200 Pro plate reader (Tecan, Männedorf, Switzerland). Exponential growth rates were calculated using the R package CellGrowth (version 3.7; Ludwig Maximilian University of Munich, [https://www.bioconductor.org/packages/release/bioc/html/cellGrowth.html]) with default settings. Growth rates were normalized to the growth rate in complete CDM.

## Results

### Manual curation produces an annotated metabolic model for S. mutans

We manually reconstructed an *in silico* metabolic model for *S. mutans* type strain UA159 (Figure 1A). Our model, named iSMU for “*in silico S. mutans*”, includes major metabolic pathways for carbohydrate metabolism and synthesis of amino acids, nucleotides, lipids, vitamins, and cofactors. The model includes 656 reactions transforming 514 metabolites. The reactions are catalyzed by the products of 488 genes (Figure 1B).

Our assembly of iSMU began with reaction databases and an annotated genome. Draft models assembled from genome annotations are often incomplete because of gaps in the genome annotation or spontaneous reactions that lack an associated enzyme. Several computational methods attempt to identify and add these missing reactions in a process called GapFilling (32). Rather than rely on automated GapFilling algorithms, we manually GapFilled iSMU by examining the reactions in each pathway. We attempted to close gaps in any pathway that 1.) was complete except for a small number of reactions, or 2.) was blocked (unable to carry flux) due to metabolites that could not be produced or consumed. We also attempted to find unannotated or misannotated genes that could catalyze the GapFilled reactions. Compared to other metabolic reconstructions, our manually GapFilled model contains fewer incomplete pathways (Table S2 and Figure 1B). On average, published metabolic models of well-studied organisms lack gene annotations for 53% of the models’ reactions (Table S2). These models also average 32.7% blocked reactions, i.e. reactions that cannot carry a steady state flux because they lack upsteam or downstream pathways. Our iSMU model has only 23.5% reactions without an associated enzyme and 2% blocked reactions.

Metabolic models simulate growth by collecting cellular building blocks into a biomass reaction. The biomass reaction is used as the objective function for metabolic simulations. A nonzero flux through the biomass reaction indicates growth in the metabolic environment specified by the model’s inputs (called exchange reactions). We modified the biomass reaction from a model of *Enterococcus faecalis* V583 (33) to create a biomass reaction for *S. mutans* UA159. Both *E. faecalis* and *S. mutans* are lactic acid bacteria with similar metabolic capabilities. To tailor the biomass reaction to *S. mutans*, we changed the relative ratios of nucleotides and amino acids. We also changed the cell membrane composition to reflect membrane sugar polysaccharides specific to *S. mutans*. We replaced UDP-N-acetyl-D-galactosamine, which based on genetic evidence is not produced by *S. mutans*, with UDP-N-acetyl-D-mannosamine and UDP-N-acetyl-D-glucosamine. We also adjusted the cell wall fatty acids to their measured proportions at pH 7.0 (34). The final biomass reaction consumes 56 metabolites to produce a unit of biomass.

### The iSMU model reveals broad metabolic capabilities of S. mutans

*S. mutans* can metabolize a wide range of carbon sources (35, 36). The ability to uptake and catabolize numerous carbohydrates allows *S. mutans* to thrive in the oral cavity of humans with varied diets. Besides glucose, the iSMU model can grow on fructose, sucrose, lactose, trehalose, ascorbate, arbutin, maltose, cellobiose, salicin, sorbitol, mannitol, mannose, N-acetyl glucosamine, fructan, galactose, galactinol, epimelibiose, melibiitol, melibiose, raffinose, maltodextrin, stachyose and malate. Growth on these carbohydrates is consistent with previous experimental studies (35, 36).

Carbohydrate metabolism is the dominant metabolic subsystem in *S. mutans* (37). Gene set enrichment for KEGG pathways classifies 134 (27.4%) of the genes in iSMU as carbohydrate metabolism (Figure 1C). By comparison, carbohydrate metabolism involves 20.7% of the genes in an *Escherichia coli* genome scale metabolic model (iJO1366, (38)) and 21.6% of the metabolic genes in a *Bacillus subtilis* genome scale model (iYO844, (39)) (Table S3).

Outside of carbohydrate metabolism, the largest subsystems in iSMU are: transport – 113 genes (23.2%); amino acid biosynthesis – 94 genes (19.3%); nucleotide biosynthesis – 56 genes (11.5%); lipid biosynthesis – 26 genes (5.3%); peptidoglycan biosynthesis – 23 genes (4.7%). (Figure 1C)

### *S. mutans* has complex nutrient requirements for growth

Several studies have investigated the minimal requirements for *S. mutans* growth *in vitro* (2, 40–43). Like many obligate human pathogens, *S. mutans* requires a combination of carbon, nitrogen, sulfur, and phosphorous sources; inorganic minerals; nucleotides; and vitamins and co-factors. We used our iSMU model and phenotypic assays to systematically explore auxotrophies for *S. mutans*.

The *S. mutans* UA159 genome encodes complete biosynthetic pathways for all 20 amino acids (37). *S. mutans* can grow without any exogenous amino acids using ammonium as the sole nitrogen source (40). The iSMU model can similarly produce biomass with ammonium and no amino acids. Using a series of leave-one-out experiments, we confirmed that the removal of individual amino acids from a defined media does not affect *S. mutans* growth in vitro (Figure 3). Simultaneous removal of cysteine and cystine does not significantly reduce growth, indicating that *S. mutans* can catabolize another sulfur source, possibly sulfate or methionine.

**Figure 3:**
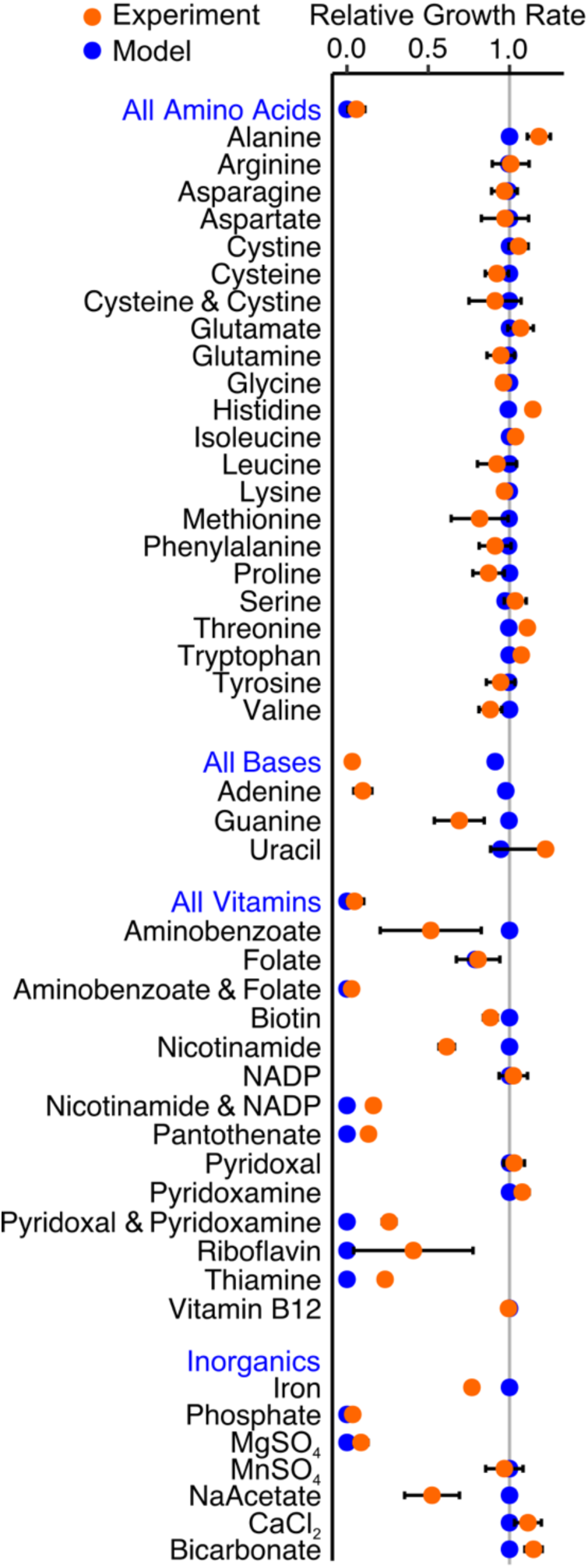
Model predictions (blue) match growth experiments for *S. mutans* UA159 in defined media (orange). Growth rate was measured for CDM lacking the specified component(s). Blue labels indicate removal of all components listed below. Experimental data are means of three independent trials with error bars representing the standard deviation. Growth rates are normalized to *S. mutans* UA159 grown in complete CDM.

*S. mutans* can theoretically synthesize nucleotides (adenine, guanine, cytosine, uracil, and thymine) but only through the non-oxidative branch of the pentose phosphate pathway (44). *S. mutans* UA159 apparently lacks the more efficient oxidative branch of the pentose phosphate pathway (45). The non-oxidative pentose phosphate pathway is bidirectional and can produce or recycle ribose 5-phosphate and other pentose sugars. These sugars are necessary precursors for nucleotide biosynthesis. The model iSMU requires no nucleotides in the media for growth. However, we found that removing all nucleotides from CDM prevents growth of *S. mutans* (Figure 3). We know that cytosine and thymine are not required for *S. mutans* growth since they are not present in CDM (Table 1). Consistent with model predictions, uracil and guanine can also be removed from CDM. Removing uracil does not significantly alter growth, but removing guanine causes a 30% decrease in growth rate (Figure 3). Only the removal of adenine completely abolished growth in CDM, which does not agree with our model predictions (Figure 3).

*S. mutans* is unable to synthesize thiamine, riboflavin, pyridoxal 5-phosphate, NAD^+^/NADP^+^, pantothenate, and folate. Anabolic pathways for these vitamins and co-factors are incomplete and key enzymes are not encoded in the *S. mutans* UA159 genome (Figure 2). All of these nutrients (or their metabolic precursors) are ingredients in two chemically defined media used to culture streptococci (CDM (31) and FMC (46)). Our model predicts aminobenzoate (an ingredient in CDM) can substitute for folate, but at least one of these nutrients is required for growth. Indeed, we found that *S. mutans* UA159 can grow in CDM without either folate or aminobenzoate but is unable to grow in media lacking both (Figure 3).

*S. mutans* cannot synthesize NAD^+^/NADP^+^ *de novo*. The iSMU model predicts that both NAD^+^ and NADP^+^ can be produced from any of NAD^+^, NADP^+^, nicotinamide, or nicotinate alone. CDM includes two of these four metabolites (NADP+ and nicotinamide). Consistent with our model, *S. mutans* can grow in CDM missing either NADP+ or nicotinamide, but not both (Figure 3).

### Gene deletion simulations match experimental data

Metabolic models contain chemical reactions and gene associations that link reactions to their corresponding enzymes. Gene associations are expressed as logical statements describing the required gene products for a reaction to carry flux. An enzymatic complex of two proteins is expressed using “and” (subunit 1 and subunit 2). A pair of isozymes that could each independently catalyze a reaction would be written with an “or” (isozyme 1 or isozyme 2). Flux balance analysis and the model’s gene associates can be combined to simulate the effects of gene deletions on growth. The logical rules in the gene associations are evaluated to identify reactions that cannot carry flux in a deletion strain. Reactions that cannot carry flux are removed from the model before calculating the maximum biomass flux. Deletion of an essential gene will prevent any nonzero biomass flux. Comparing experimentally determined essential genes with the model’s predictions to validate the model’s gene associations.

We simulated the effects of all single deletions for the 488 genes in the iSMU model. We compared the *in silico* deletions to two experimental gene deletion studies in *S. mutans* UA159: a transposon mutagenesis sequencing (Tn-seq) experiment (47) and a screen of an ordered array of single gene deletion strains (48). The Tn-seq study used a mariner-family transposon to generate random insertions across the UA159 genome (47). The transposon/genomic DNA junctions were amplified and sequenced to quantify fitness after growth in a defined media (FMC). Genes lacking transposon insertions sites are predicted to be essential in FMC. Overall, 84.8% of the essentiality predictions from the iSMU model were consistent with the Tn-seq data (Figure 4B). A comparison between iSMU’s essential gene predictions and the Tn-seq data is shown in the iSMU map in Figure 2.

**Figure 4:**
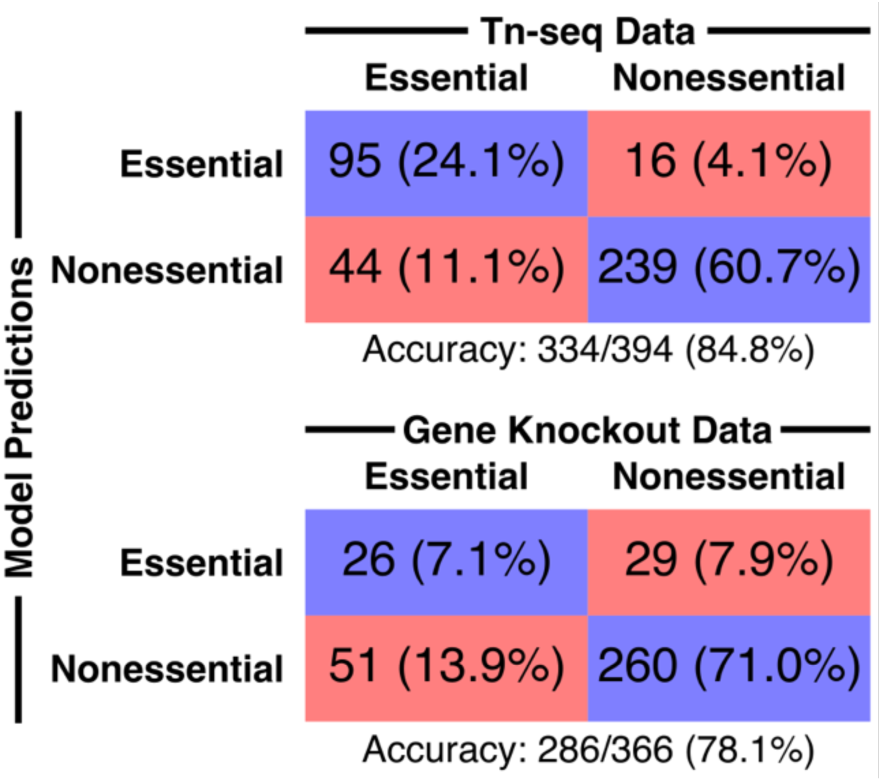
iSMU essentiality predictions align with two experimental studies. Shields et al. (47) (top) used transposon mutagenesis sequencing (Tn-seq) to identify essential genes in the defined media FMC. Quivey, et al. (48) (bottom) screened a library of single gene knockout strains for growth in rich media. Blue boxes indicate the number (percentage) of genes in the model and dataset that are both essential or nonessential. Red boxes indicated disagreements between the model and experiments.

The gene deletion strains in Quivey, et al., (48) were constructed individually using homologous recombination with a selective marker. The deletion library contains strains for 1112 of the 1956 genes in *S. mutans* UA159, including 366 of the 488 genes in the iSMU metabolic model. The remaining 122 genes are hypothesized to be essential or could not be deleted due to technical limitations. Deletion strains were grown in Brain Heart Infusion media (BHI), a rich and undefined media. We simulated BHI by opening all exchange reactions in the iSMU model. As shown in Figure 4A, 78.1% of the experimental essentiality results agreed with the model predictions. Agreement between iSMU’s essential gene predictions and the data from Quivey, et. al. is highlighted the iSMU map in Figure S1.

### Manual curation improves model consistency and accuracy

Several software systems can generate draft metabolic models using reaction databases and annotated genomes. Our first attempt at reconstructing the metabolism of *S. mutans* used a draft model from the KBase system (49). Unfortunately, the draft model lacked many of the metabolic features of lactic acid bacteria and included several subsystems known to be inactive in homofermentive anaerobes. We therefore abandoned the KBase model and began a manual reconstruction process. We compared the final iSMU model to the KBase model to quantify the disagreement between the manual and automated reconstruction pipelines.

Our manually curated reconstruction differs substantially from the reconstruction produced automatically by KBase. Our iSMU model has 6.4% fewer reactions than the KBase model (656 vs. 701) but 97% more genes (488 vs. 247) (Figure 1B). Thus, the manually curated model has a larger proportion of gene associated reactions than the automated reconstruction. The dearth of gene associations in the KBase model is due in part to the 237 reactions added during GapFilling, since GapFilled reactions are added without genomic evidence for the reaction. By comparison, our iSMU model required only 11 GapFilled reactions to enable growth on defined media and 19 carbon sources.

The KBase model contains 41% more metabolites than our iSMU model. Unfortunately, 22 of the metabolites in the KBase model are “dead-end” metabolites that lack either a producing or consuming reaction. The dead-end metabolites block flux through 299 (43%) of the KBase model’s reactions. At steady-state, these blocked reactions cannot carry flux or be analyzed using Flux Balance Analysis. Less than 2% of the reactions in our iSMU model are blocked, indicating more complete reaction pathways than the automated reconstruction.

## Discussion

iSMU is the first whole genome metabolic model of the cariogenic pathogen *S. mutans* UA159. The model captures the entire metabolism of the organism and was validated by comparing model predictions to experimental evidence. Metabolism plays a dual role in the pathogenicity of *S. mutans*. First, fermenting sugars creates caries-causing lactic acid. A significant portion of the *S. mutans* genome is dedicated to carbohydrate metabolism, reflecting the plasticity of *S. mutans*’ metabolism. Second, multiple metabolic subsystems are required for *S. mutans* to tolerate acid and outcompete non-cariogenic streptococci. A mathematical model allows us to investigate connections among metabolic pathways during pathogenesis.

iSMU’s predictions agree with most of the nutrient depletion experiments, but some of *S. mutans*’ auxotrophies are unexplained by the model. For example, the UA159 genome encodes a complete pathway for adenine synthesis, but exogenous adenine is required for growth *in vitro*. The adenine synthesis pathway in iSMU may not be expressed or functional in UA159 when grown aerobically in CDM. Other experiments agree qualitatively, but not quantitatively with the model. When guanine, aminobenzoate, nicotinamide, or sodium acetate are removed from CDM, S. mutans grows slower predicted by the model. The model also underpredicts growth rates when pyridoxal and pyridoxamine, riboflavin, and thamine are removed. Differences like these are expected with constraint-based models that lack kinetic details for nutrient uptake and enzymatic turnover.

*S. mutans* UA159 can grow on minimal media with ammonium as the sole nitrogen source (40), and the iSMU model can produce biomass in these conditions. Experimentally, growth on ammonium requires an anaerobic environment, but the model can produce biomass with or without oxygen. Oxygen may repress expression of enzymes required for scavenging nitrogen from ammonium, and the lack of regulation in our model would explain why iSMU can grow aerobically using ammonium.

Several factors could explain the disagreements between the model’s essentiality prediction and experimental results. First, we note that both methods for identifying essential genes have technical strengths and limitations. In the ordered gene deletion library, any gene for which a deletion mutant cannot be constructed is labeled essential. A nonessential gene located in a region of the chromosome refractory to homologous recombination would be incorrectly labeled as essential. The Tn-seq libraries in Shields, et al. were constructed using *in vitro* transposon mutagenesis followed by homologous recombination, so the same limitation applies. The Tn-seq libraries were grown for ∼30 generations before sequencing. Such a large expansion can bias the library against mutants with large fitness defects. Although these mutants may be viable, they appear at such low frequency in the final pool that they are missed during sequencing. The corresponding genes would be incorrectly labeled as essential. The disagreement between the Tn-seq and defined deletion library suggest the “essential genome” of *S. mutans* UA159 has not been fully elucidated.

Inaccuracies in the *iSMU* model also contribute to disagreements over essential genes. After decades of curation, metabolic models for the model organisms *E. coli* and *S. cerevisiae* still miss some essential gene predictions (38, 50). Unannotated genes could catalyze redundant routes to synthesize essential metabolites *in vivo*, creating false positive essentiality predictions in iSMU. Regulation, loss of function mutations, and missing cofactors can also restrict the metabolic capabilities of *S. mutans*, making the pathogen less metabolically flexible than the iSMU model.

Overall, believe the model’s predictions could be improved by either 1.) incorporating regulatory rules during simulations, or 2.) using gene expression or other high-throughput data to tailor the model to anaerobic, aerobic, acidic, or other conditions. *S. mutans*’ metabolic requirements change as the bacterium encounters different niches in the mouth. Before forming thick biofilms and deep dental caries, growth conditions are likely aerobic with abundant nutrients from saliva and food consumed by the host. Deep dental caries may create anaerobic conditions with limited nutrient availability. In this environment, *S. mutans* would need to synthesize many biomass components de novo.

*S. mutans* is a model organism in oral microbiology (51). Our iSMU model draws from hundreds of studies to form an accurate, genome-wide picture of *S. mutans* metabolism. The model also highlights the value added by manual curation. The metabolism of *S. mutans* is well characterized on the molecular and pathway levels. Incorporating manually curated models of *S. mutans* and other lactic acid bacteria may improve the accuracy of automatic reconstruction pipelines.

## Acknowledgements

We thank Robert Shields for assistance with the Tn-seq data and John Gerlt for sharing equipment and reagents. This work was supported by the NIH National Institute of Dental and Craniofacial Research (grant DE026817 to PAJ) and the University of Illinois at Urbana-Champaign. The authors declare no financial conflict of interest.

## Supplemental Material

**Table S1**: GapFilled reactions in iSMU and evidence for inclusion.

**Table S2**: Curated genome-scale metabolic models used to compare number of reactions without a gene association and blocked reactions.

**Table S3**: Model comparisons between iSMU, iJO1366, and iYO844.

**File S1**: iSMU genome-scale metabolic model in SBML file format.

**File S2**: iSMU genome-scale metabolic model in spreadsheet file format.

**File S3**: iSMU’s entire model map in JSON file format.

**File S4**: iSMU’s entire model map in SVG file format.

